# Repeatable differences in exploratory behaviour predict tick infestation probability in wild great tits

**DOI:** 10.1101/2020.03.06.978973

**Authors:** Robert E. Rollins, Alexia Mouchet, Gabriele Margos, Volker Fingerle, Noémie S. Becker, Niels J. Dingemanse

## Abstract

Ticks are parasites that feed on the blood of various vertebrate hosts, including many species of bird. Birds can disperse ticks over short and long distances, therefore impacting tick population dynamics. The likelihood that birds attract ticks should depend on their behaviour and the environment. We studied various key ecological variables (breeding density, human disturbance) and phenotypic traits (exploratory behaviour; body condition) proposed to predict tick burden in great tits (*Parus major*). Our study spanned over three years and 12 human-recreated plots, equipped with nest-boxes in southern Germany. Adult breeders were assessed for exploratory behaviour, tick burden, and body condition. For each plot, human disturbance was quantified as a human recreational pressure index during biweekly nest box inspections by scoring the number of recreants using the plots. Infestation probability but not tick burden increased with exploratory behaviour. We also found moderate support for a positive effect of recreational pressure on infestation probability. Further, body condition negatively predicted tick burden. Individuals were repeatable in tick burden across years. Our study implies that infestation probability and tick burden are governed by distinct ecological and phenotypic drivers. Our findings also highlight the importance of incorporating ecological and individual variation in host phenotypes to predict spatiotemporal distributions of ticks in nature. (207/250-word limit)

**Lay Summary:** Ticks use many birds as hosts, but why do some individuals have more or fewer ticks? Using a data collected over three years on great tit adults inhabiting 12 different nest-box plots, we showed that more explorative birds and those in highly recreated habitats were more likely to be infested with ticks. Exploratory behaviour and human disturbance could modify great tit habitat choice and, therefore, impact how often a birds and ticks encounter each other.

## 1. Introduction

Ticks are obligate ectoparasites and require a blood-meal between each distinct life stage (Hillyard 1996; Kurtenbach et al. 2006).Thus, tick population dynamics depend on host movement and behaviour (Hillyard 1996; Hasle 2013). In turn, ticks can influence host population dynamics by affecting host fitness (Heylen et al. 2009), condition (Heylen and Matthysen 2008; Müller et al. 2013; Norte et al. 2013), and behaviour (Hoodless et al. 2002; Barber and Dingemanse 2010; Sih et al. 2018) through blood feeding and potential pathogen transfer (Stanek 2009; Norte et al. 2013). Bird species are important hosts of various tick species, disperse ticks over short and large distances (Hasle 2013), and thus affect tick-borne pathogen dynamics (Stanek 2009; Hasle 2013). Insight into the main sources of variation in avian tick uptake and burden is thus required to predict the spatiotemporal distribution of ticks in nature.

The likelihood and number of ticks carried by an individual bird vary as a function of multiple interacting ecological factors affecting tick survival in the breeding habitat, such as temperature or humidity (Lindgren et al. 2000; Oorebeek and Kleindorfer 2008; Kiffner et al. 2011; Titcomb et al. 2017), host breeding density (Oorebeek and Kleindorfer 2008; Takumi et al. 2019), and vegetation type/coverage (Kiffner et al. 2011; К. Tack et al. 2012; W. Tack et al. 2012; Tack et al. 2013), and are thus predicted to covary with habitat modification and climatic change (Lindgren et al. 2000; Randolph 2004; Akimov and Nebogatkin 2016). For example, city-dwelling birds may carry lower tick burdens (Evans et al. 2009; Hamer et al. 2012) as ticks are less abundant in urban environments (Akimov and Nebogatkin 2016). However, birds in worse physiological condition may also attract more ticks because they are more susceptible to take up parasites (Barber and Dingemanse 2010; Abbey-Lee et al. 2016; Hutfluss and Dingemanse 2019). Further studies documenting ecological predictors of tick burden are therefore required to firmly establish factors with general versus study-specific predictive effects (Kelly 2006; Nakagawa and Parker 2015). In this study, we quantified tick burdens on an avian host, the great tit (*Parus major*), within 12 nest box plots surveyed for breeding density and human disturbance levels across three breeding seasons. This enabled us to investigate the role of these two key environmental factors in shaping avian tick burden.

Tick burden can also vary among individual birds as a function of individual-specific traits, such as behaviour (Barber and Dingemanse 2010; Newman et al. 2015; Sih et al. 2018) or body condition (Norte et al. 2013; Newman et al. 2015). For instance, bird species with larger average body sizes have increased tick burdens compared to smaller bird species (Newman et al. 2015) as do bird species with a higher proportion of ground foraging behaviour (Newman et al. 2015; Loss et al. 2016; Kocianová et al. 2017). Similarly, within populations of the same species, individual-specific differences in foraging or social behaviour can also predict tick burdens (Sih et al. 2018). For example, individuals that are more active and explorative in their (foraging) behaviour may encounter more parasites (i.e. ticks), and consequently recruit higher tick burdens (Barber and Dingemanse 2010; Sih et al. 2018; Taggart et al. 2018). Body condition and exploratory behaviour are both repeatable traits in our great tit populations (Stuber et al. 2013; Moiron et al. 2018; Moiron et al. 2019). To quantify effects of body condition and exploratory behaviour on tick burden in this study, we assayed both traits on all birds in each year, such that effects on tick burden could be estimated among-individuals (Niemelä and Dingemanse). Among-individual relationships imply that long-term repeatable differences in phenotypic traits predict tick burden. Sex and age may also play a role in avian tick burden as males (versus females) tend to have higher tick burdens and older (versus younger) birds have lower tick burdens (Heylen and Matthysen 2008; Heylen et al. 2013).

The overall aim of this study was to test effects of environmental factors (breeding density and human disturbance) and phenotypic traits (body condition, exploratory behaviour) on tick burden in the wild. Our repeated measures design additionally enabled the quantification of the amount of variance in tick burden attributable to among-year (temporal variation), among-plot (spatial variation), and among-bird (individual variation) variation, thereby acquiring quantitative estimates of the magnitude of variation attributable to multiple biological levels of variation.

## 2. Material and Methods

### 2.1. Study sites and data collection

Our study was performed on 12 nest box plots within a 10×15 km^2^ area in southern Munich (47° 97’N, 11° 21’E) which were established in autumn 2009. Each plot was fitted with 50 nest boxes in a regular grid covering approximately 9 hectares. Breeding parameters (detailed below) were monitored for three years (2017-2019) within all plots except one plot that was monitored only in 2017. Each nest-box was inspected biweekly during the breeding seasons (April-July) for nesting activity. During such plot inspection, the number of recreants was counted as a measure of human disturbance (detailed by Hutfluss & Dingemanse, (2019)).

When hatchlings were 10-12 days old, both parents were captured in the nest box using a spring trap. Un-banded birds were given a unique, numbered band. Birds were then tested for their exploratory behaviour (detailed below), weighed, morphologically measured (Moiron et al. 2019), aged based on plumage characteristic as first year breeder or older (Dingemanse et al. 2020), and assessed for ticks (protocol detailed below). All birds were captured in May (with a few captures in June in 2018).

### 2.2. Exploratory behaviour

Exploratory behaviour was measured using a novel environment test adapted to the field from established protocols (Stuber et al. 2013). Birds were transferred to a holding box attached to a cage (61L × 39W × 40 H cm^3^) fitted with a mesh front and three perches, representing the novel environment. The focal bird was released into the novel environment without handling and recorded for two minutes with the field observer located out of sight. Videos were subsequently scored by dividing the cage into six equal sections and three floor sections as described in (Stuber et al. 2013). Exploration score was calculated as the sum of movements the individual did between these sections (Abbey-Lee and Dingemanse 2019; Dingemanse et al. 2020).

### 2.3. Tick burden and collection

Ticks tend to concentrate around the eyes and beak of great tits (Heylen and Matthysen 2008). A regional patch examination protocol (reviewed in (Lydecker et al. 2019)) was developed to standardize the screening: 1) screen around the eyes/ears and along the margin of the beak on both sides of the bird, 2) screen underneath the beak, 3) screen along the top margin of the beak, 4) screen the top of the head. All captured birds were screened in 2018-2019, while in 2017 only a sample of birds were screened instead. The total number of ticks carried by each captured bird was recorded. In 2018 and 2019, ticks were collected using fine tweezers and stored in 99% ethanol as part of another study.

### 2.4. Recreational pressure index

During each plot inspection, all observers recorded each recreant seen and its specific location (see (Hutfluss and Dingemanse 2019)). Each recreant was connected to the first observation to avoid double counts. The probability to observe a recreant during a plot inspection is biased by various factors (Hutfluss and Dingemanse 2019). To obtain an unbiased index of recreational pressure, the binary probability to observe recreants was calculated using all inspections conducted 2010-2019 (n = 3724 inspections). This probability was calculated using a binomial generalized linear mixed effects model (GLMM) fitting fixed effects for plot inspection duration, the number of observers, and starting time (in hours from sunrise) (Hutfluss and Dingemanse 2019) We fitted random intercepts for each unique combination of plot and year (termed plot-year, see (Abbey-Lee et al. 2016; Araya-Ajoy et al. 2016; Araya-Ajoy and Dingemanse 2017)) to acquire an average value of recreational pressure for each plot in each year, and for date of observation to control for date-specific environmental effects. We extracted best linear unbiased predictors (BLUPs) for each plot-year between 2017-2019 (n = 34) and used them in subsequent models as a recreational pressure index. The usage of BLUPs has been criticised when associated uncertainty is not taken forward; some have proposed to estimate a posterior distribution of possible BLUP values and thereby take forward uncertainty in subsequent analyses (Hadfield et al. 2010; Houslay and Wilson 2017). However, recent work has shown that taking forward uncertainty in BLUP-values resulted in biased estimates, whereas utilizing average BLUPs as fixed effects was shown to be less precise but unbiased (Dingemanse et al. 2020). Therefore, we here present the estimated effect of the average BLUP values.

### 2.5. Data Preparation

The scaled mass index (Peig and Green 2009) was used as our measure of body condition. Briefly, each body mass measurement was multiplied by the mean tarsus length of the population (separately for each sex) which was first divided by the individual tarsus length (measured at the same time as body mass) raised to a scaling exponent calculated from a log transformed regression of tarsus on body mass measurements as described in Peig & Green, (2009). This regression was calculated using all bird records from 2010-2019 (n = 4273 records) which included individuals repeatedly captured and measured.

Certain factors, such as experience (Dingemanse et al. 2012; Dingemanse et al. 2020), influence exploration scores and need to be corrected for to avoid biased measures. Individual mean exploratory scores were calculated using a linear mixed effects model (LMM) on exploratory behaviour data from all years (2010-2019) (n = 4251) (outlined in (Dingemanse et al. 2020)) assuming a Gaussian error distribution. Sequence number (i.e. the number assays per bird) was fitted as a fixed effect covariate, as well as individual and plot-year as random intercepts. An average BLUP per individual, representing an exploratory score corrected for experience, was extracted and used in subsequent analysis as a measure of an individual’s average exploratory score.

Density was calculated as the number of breeding pairs producing first clutches per hectare.

### 2.6. Statistical Analysis

Tick infestation of great tits was analysed in two parts using GLMMs. First, infestation probability was estimated using all recorded captures between 2017-2019 (n = 784 records). Second, tick burden (i.e. absolute number of ticks observed) was modelled using only infested individuals (n = 520 records). For both response variables, the model set up was the same. Fixed effects were fitted for recreational pressure, breeding density, body condition, sex, age, and mean individual exploration score. Random intercepts were fitted for year, plot, plot-year, field observer, nest-box, and individual. Prior to analysis, all fixed effects were variance standardized, such that the statistical intercepts of our models reflected the value for the average individual in the average environmental condition.

All statistical analyses were performed in R (version 3.5.3) (R Core Team 2019). Models were run using the functions *lmer* and *glmer* from the package *lme4* (Bates et al. 2015) assuming either binomial (infestation probability) or Poisson (tick burden) error distributions. Mean estimates and their 95% credible intervals (CI) were estimated based on 5,000 simulations using the *sim* function from the *arm* package (Gelman and Su 2016). Residual errors were calculated according to Nakagawa & Schielzeth, (2010). Adjusted repeatability values were calculated as the proportion of variance unexplained by the fixed effects that was explained by the focal random effect. Effects of fixed effects were considered “strongly supported” when their 95% CI did not overlap zero or “moderately supported” if the point estimate was skewed away from but overlapped zero. Estimates that were centred on zero were viewed as having strong support for the absence of an effect (Dingemanse et al. 2020).

## 3. Results

Adult great tits carrying ticks were found in all study plots and all years (Figure 1) with an average tick burden of (mean ± s.e.) 5.63 ± 0.33 ticks per infested bird (Figure 2). Infestation probability varied between plots (Table 1) but not tick burden (Table 1). Both tick burden and infestation probability, varied among nest-boxes (Table 1). Individuals were not repeatable in infestation probability (Table 1) but were moderately repeatable in tick burden (mean parameter estimate: 0.44; 95% CI: (0.39; 0.50)). Exploratory behaviour positively predicted infestation probability (0.18 (0.01; 0.35), Figure 3) but not tick burden (Table 1). Additionally, great tit body condition negatively predicted tick burden (−0.11 (−0.19; −0.02), Figure 4) but not infestation probability (Table 1). Recreational pressure positively predicted infestation probability with moderate support. (0.21 (−0.03; 0.43), Figure 5). Great tit breeding density, age, and sex did not predict infestation probability or tick burden (Table 1).

**Table 1.** Effect size estimates (β) and 95% credible intervals (CI) for great tit tick infestation as the probability of infestation (n = 784 records) and tick burden of infested individuals (n = 520 records).

|  | Infestation Probability | Tick Burden |
| --- | --- | --- |
| **Fixed Effects** | **$\beta$ (95% CI)** | **$\beta$ (95% CI)** |
| Intercept | 0.89 (0.42, 1.37) | 1.20 (0.75, 1.64) |
| Recreational pressure | 0.21 (-0.03, 0.43) | -0.07 (-0.22, 0.08) |
| Breeding density | 0.08 (-0.26, 0.43) | -0.04 (-0.25, 0.17) |
| Exploratory behaviour | 0.18 (0.01, 0.35) | -0.02 (-0.12, 0.09) |
| Body condition | 0.06 (-0.11, 0.24) | -0.11 (-0.19, -0.02) |
| Sex<sup>a</sup> | -0.06 (-0.42, 0.24) | -0.06 (-0.24, 0.12) |
| Age | -0.07 (-0.24, 0.09) | 0.01 (-0.08, 0.10) |
| **Random Effects** | **$\sigma^2$ (95% CI)** | **$\sigma^2$ (95% CI)** |
| Year | 0.04 (0.00, 0.17) | 0.13 (0.03, 0.36) |
| Plot | 0.47 (0.18, 0.94) | 0.05 (0.02, 0.09) |
| Plot-Year | 0.08 (0.04, 0.12) | 0.07 (0.04, 0.12) |
| Observer | 0.01 (0.00, 0.01) | 0.13 (0.06, 0.23) |
| Nest-box | 0.64 (0.55, 0.74) | 0.28 (0.23, 0.32) |
| Individual | 0.00 (0.00, 0.00) | 0.45 (0.40, 0.51) |
| Residual | 0.35$\pi^2$ (0.35$\pi^2$, 0.34$\pi^2$) | 1.27 (1.39, 1.18) |
| **Adjusted Repeatability** | **$r$ (95% CI)** | **$r$ (95% CI)** |
| Year | 0.01 (0.00, 0.03) | 0.05 (0.01, 0.13) |
| Plot | 0.10 (0.04, 0.18) | 0.02 (0.01, 0.03) |
| Plot-Year | 0.02 (0.01, 0.02) | 0.03 (0.02, 0.04) |
| Observer | 0.00 (0.00, 0.00) | 0.05 (0.03, 0.08) |
| Nest-box | 0.14 (0.13, 0.14) | 0.12 (0.11, 0.11) |
| Individual | 0.00 (0.00, 0.00) | 0.19 (0.18, 0.18) |
| Residual | 0.74 (0.82, 0.63) | 0.53 (0.64, 0.42) |

<sup>a</sup>Differences between sexes (reference category = female)

**Figure 1.**
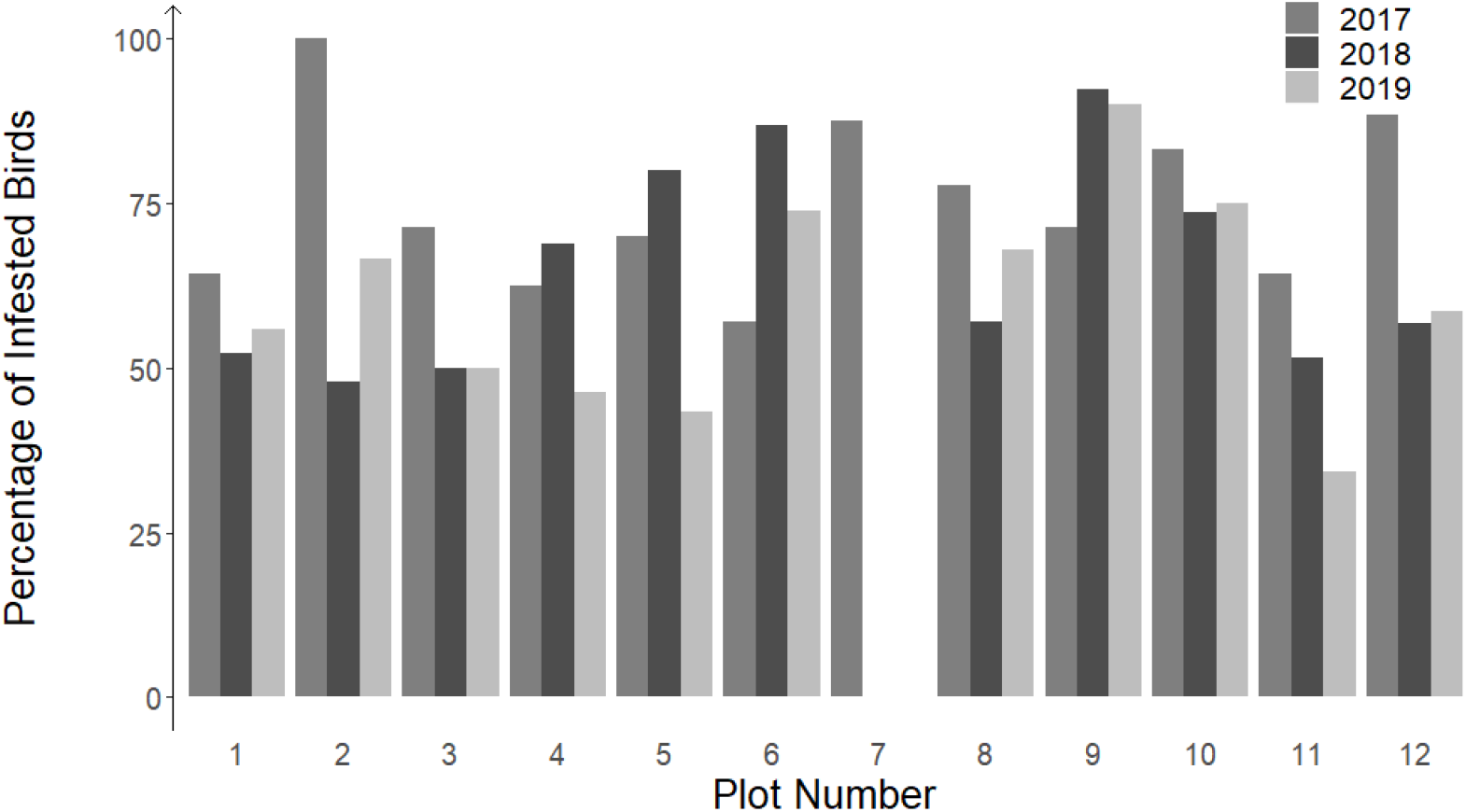
Percentages of all captured great tit adults infested with ticks from the 2017-2019 breeding seasons. Data for plot 7 were only available in 2017.

**Figure 2.**
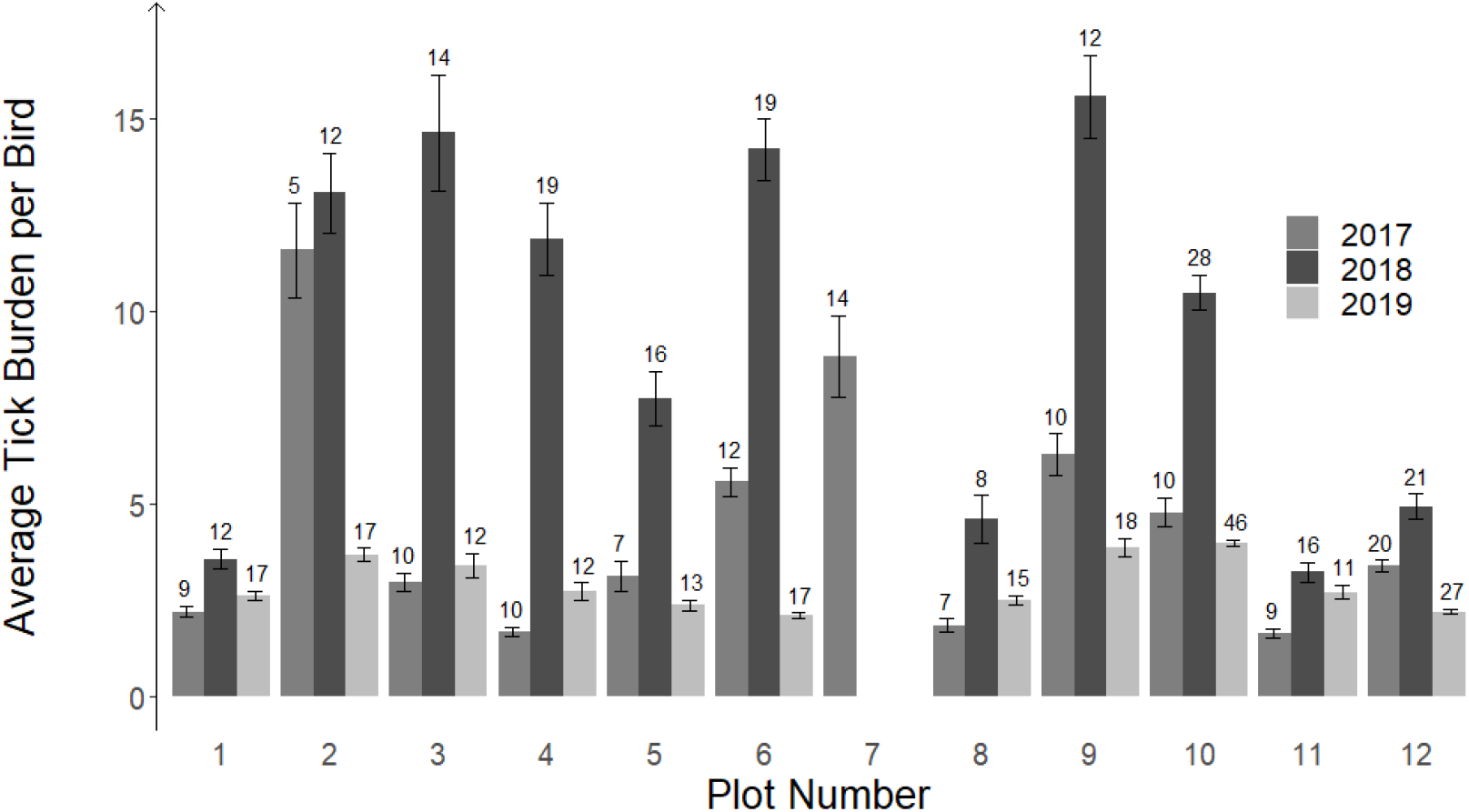
Average tick burden per infested great tit adult captured in each study plot during each field season (years 2017-2019). The error bars represent standard error and the numbers above each bar reflect the absolute number of infested great tit adults per plot. Data for plot 7 were only available in 2017.

**Figure 3.**
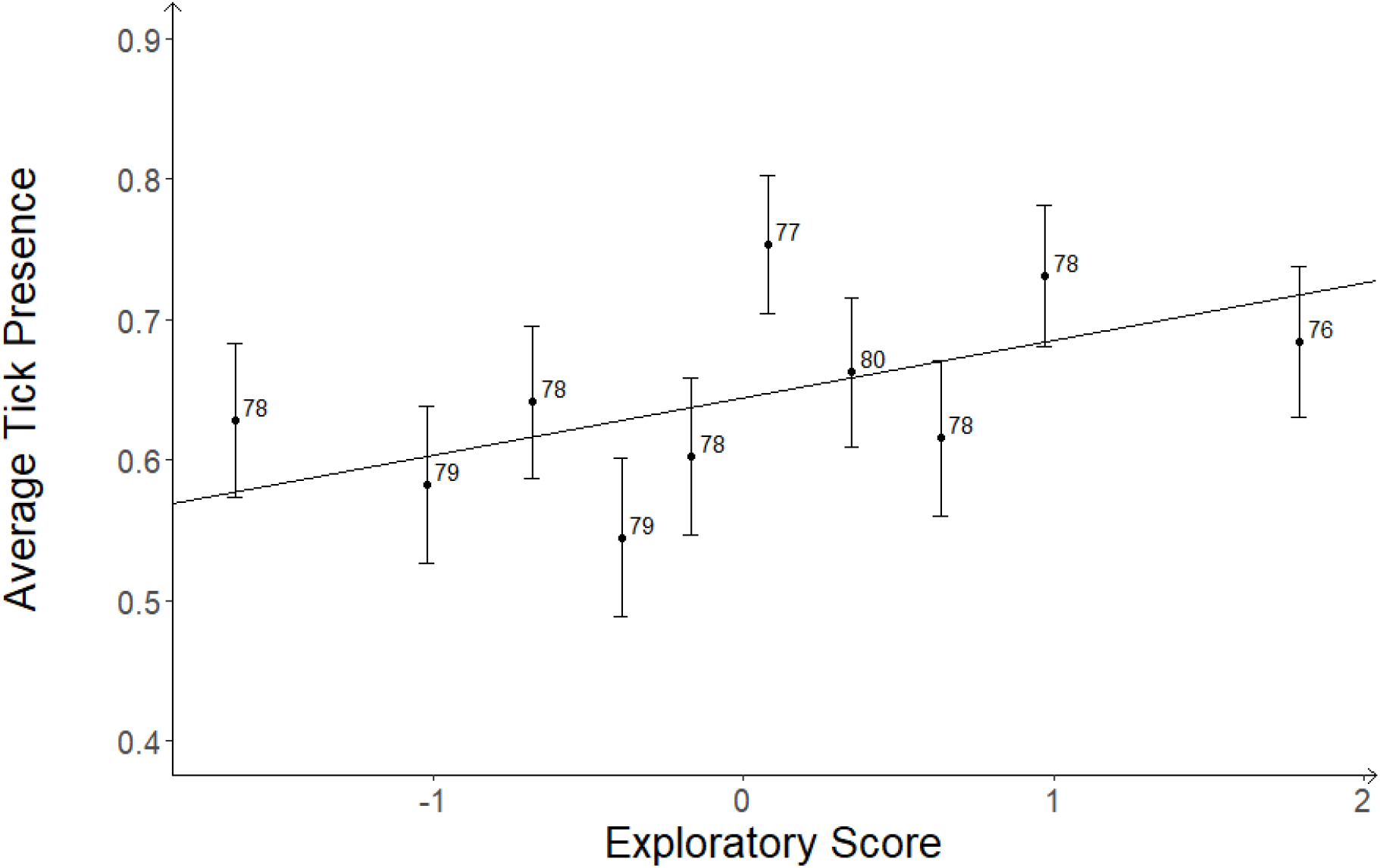
Tick presence in relation to exploratory score. For graphical purposes only, data were split into 10 quantiles and were averaged to better show the linear relationship. Each point refers to the mean tick presence and exploratory score for that quantile including data from all collection years (2017-2019). Error bars report the standard error for the respective mean tick presence. The numbers report the number of records in the respective bin. The line reports the linear relationship from the model.

**Figure 4.**
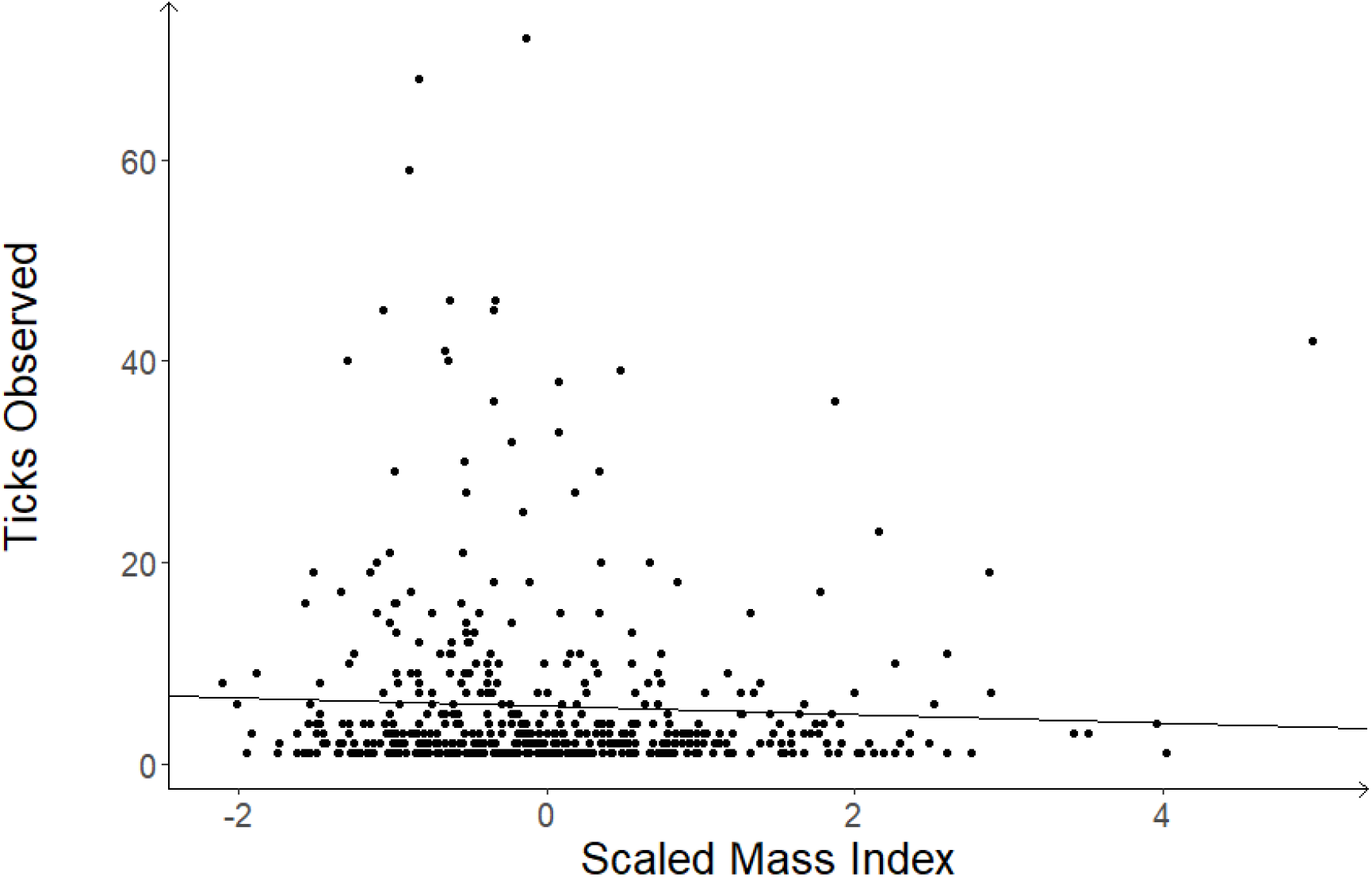
Tick burden of infested birds in relation to birds scaled mass index (i.e. body condition). Each point refers to an individual infested bird record. Records from all collection years (2017-2019) are included. The line reports the linear relationship from the model.

**Figure 5.**
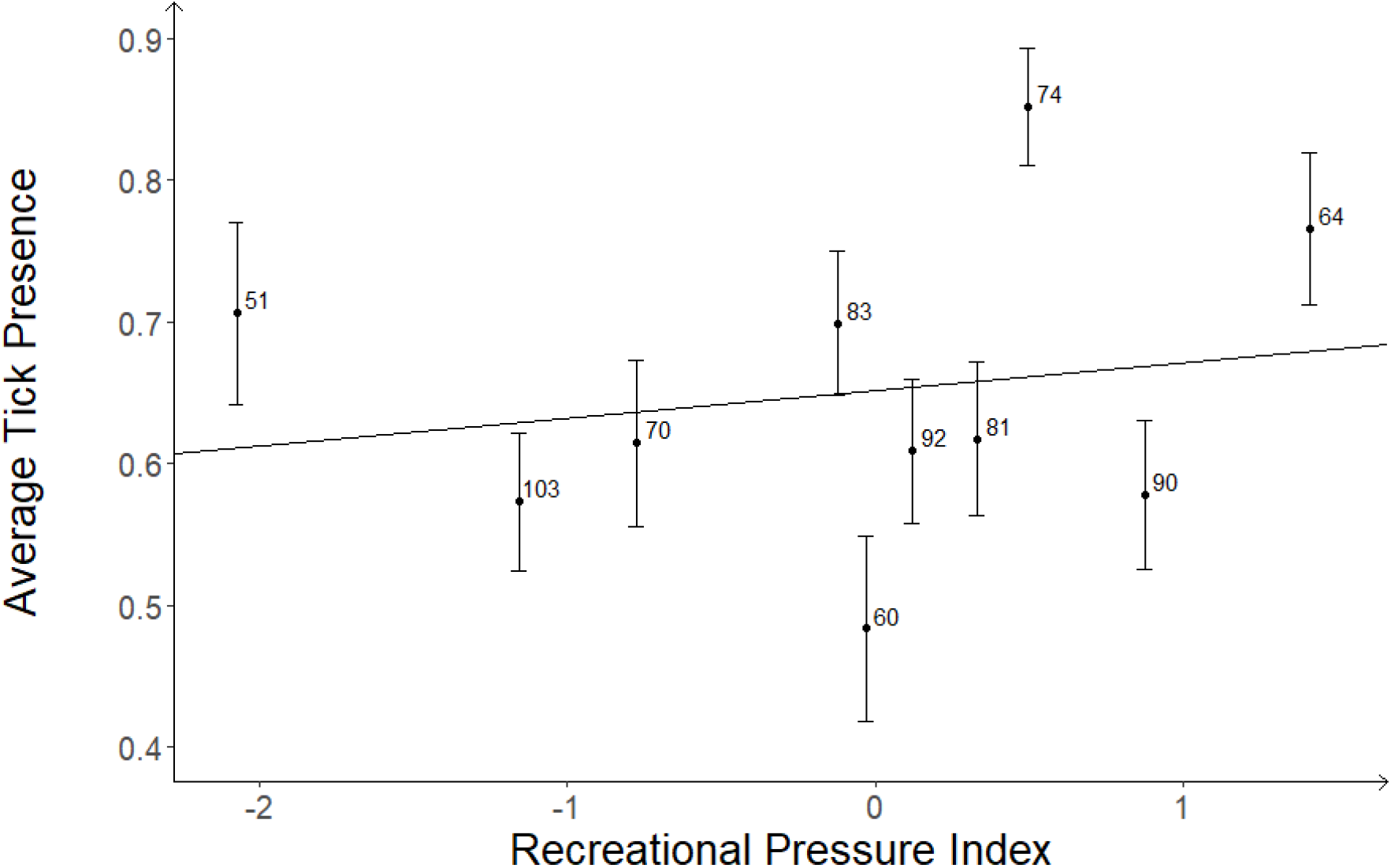
Tick presence in relation to recreational pressure index. For graphical purposes only, data were split into 10 quantiles and were averaged to better show the linear relationship. Each point refers to the mean tick presence and recreational pressure index for that quantile including data from all collection years (2017-2019). Error bars report the standard error for the respective mean tick presence. The numbers report the number of records in the respective bin. The line reports the linear relationship from the model.

## 4. Discussion

Birds influence the spread of tick species, thus to understand tick population dynamics, the underlying drivers of avian tick infestation need to be determined (Hasle 2013; Heylen et al. 2013). Here we quantified effects of proposed environmental and phenotypic predictors on tick infestation (Newman et al. 2015) in the great tit. More explorative birds and those living in highly recreated plots had increased infestation probabilities, while birds with lower body condition had higher tick burdens. Both tick burden and infestation probability varied across time (years) and space (plots). Individual tick burden was moderately repeatable across years, implying that individual-specific phenotypic traits not studied here influence tick infestation in great tits.

Some of the variation in tick infestation could be due to habitat choice of the great tits. Several ecological factors might influence in which habitats those birds nest, such as resource availability, conspecific density (Spiegel et al. 2015; Spiegel et al. 2017; Sih et al. 2018), or human disturbance (Hutfluss and Dingemanse 2019). Great tit population density and human disturbance might push some individuals to occupy habitats more suitable for ticks, e.g. more humid habitats with denser vegetation (К. Tack et al. 2012; W. Tack et al. 2012), therefore influencing tick infestation. Breeding density did not influence tick infestation; however, human disturbance might have as tick infestation was higher in plots with higher recreational pressure and varied among nest-boxes within plots. A previous study showed that, great tits preferentially nested further away from heavily recreated paths (Hutfluss and Dingemanse 2019) supporting the fact that recreational pressure can modify habitat choice. Recreational pressure however, as breeding density, did not predict tick burden. Tick burden did vary among plots and nest-boxes implying that other environmental factors influence great tit tick burden. For example, several ecological factors, such as relative humidity (Lindgren et al. 2000; Oorebeek and Kleindorfer 2008; Kiffner et al. 2011), vegetation (Kiffner et al. 2011; W. Tack et al. 2012; К. Tack et al. 2012; Tack et al. 2013), or host availability (i.e. rodents, roe deer, etc) (Oorebeek and Kleindorfer 2008; Kiffner et al. 2011; Takumi et al. 2019) might impact tick survival and distribution, leading to spatial heterogeneity in tick abundance and, consequently, tick burden on hosts.

Various phenotypic traits varying among individual birds might also influence tick infestation. Tick infestation probability, but not tick burden, increased with individual exploratory behaviour. This pattern has previously been observed in Eastern grey squirrels (*Sciurus carolinensis*) (Santicchia et al. 2019). Exploratory behaviour is known to predict foraging style (Verbeek et al. 1994; Aplin et al. 2014) but to which extent is not fully understood (Moyers et al. 2018). *Ixodes ricinus* (the most common tick infesting vertebrates in Europe (Hillyard 1996)) quests for hosts vertically up to 1.5m but with decreasing abundance with increasing distance from the ground (Mejlon and Jaenson 1997). More explorative birds might forage more within this questing range and closer to the ground and, therefore, increase their encounter rate with ticks. Other research has argued that other parasite burdens than tick burden should increase with body mass instead of exploration (Santicchia et al. 2019). Some studies have indeed found that parasite burdens can increase with body mass (Kiffner et al. 2011; Kiffner et al. 2013; Mysterud et al. 2015). However, this relationship has not been observed in great tits (Heylen and Matthysen 2008; Heylen et al. 2013) and other commonly studied species (Kiffner et al. 2013). Moreover, effects of both exploratory behaviour and body mass have not been simultaneously investigated and thus not disentangled. By jointly fitting exploratory behaviour and body condition in our analysis we could properly differentiate between these two effects. Previously, the absence of a relationship between great tit body mass and tick burden has been supported (Heylen and Matthysen 2008). Our study partially confirms this result as body condition did not predict infestation probability, though it negatively predicted tick burden. This negative relationship between body condition and tick burden has been observed in other species and has been attributed to health reductions due to blood loss (Norte et al. 2013). Both age and sex did not influence tick infestation in our study. Previous literature showed that males tend to have increased tick burdens (Hoodless et al. 2003) and that tick burden decreases with age (Heylen et al. 2013). Other work however showed no effect of age on tick burden as in our study (Heylen and Matthysen 2008). The null effect of sex though could be due to the time of data collection where adults might be in a lower condition due to chick rearing (Ots and Horak 1996), and thus any sex differences might be masked. Infested individuals were, however, repeatable for their tick burdens meaning that other phenotypic traits than those measured here (i.e. exploratory behaviour and body condition) might explain tick burden of infested birds. This could be other behaviours such as aggression (Zohdy et al. 2017) or foraging behaviour (Newman et al. 2015; Loss et al. 2016; Kocianová et al. 2017) as both play roles in tick burden.

It is important to mention that birds were captured approximately over a one-month period (range: 25-29 days) mostly during May each year which corresponds to the peak of tick abundance (Kurtenbach et al. 2006). However, some variation in tick infestation could be due to weather related conditions when the birds encountered the ticks. Weather and climatic variables affect questing activity of ticks (Mejlon and Jaenson 1997) and consequently the probability for a bird to encounter a tick and ultimately overall tick infestation. Additionally, both breeding density and bird age where shown to have null effects on tick infestation. However, this null result could be due to effects at different levels of biological variation being confounded (van de Pol and Wright 2009). To test if this was the case, we split age into among- and within-individual variation in age and split breeding density into temporal- and spatial-variation in density. These level-specific analyses also failed to show effects of these factors (Supplemental Text 1) supporting the absence of these effects.

## 5. Conclusions

We showed that exploratory behaviour predicted tick infestation probability in great tits, with more explorative birds having an increased infestation probability. However, tick burden of infested birds was not influenced by exploratory behaviour, sex, or age but was negatively predicted by body condition. We additionally showed that individuals were repeatable for tick burden, meaning that other phenotypic traits not analysed here might explain consistent differences in tick burden of infested individuals. We also found moderate support for a positive effect of recreational pressure on infestation probability. Our results highlight the complex nature of tick infestation and how various factors influence infestation probability and tick burden.

## 6. Ethics Statement

All procedures complied with guidelines from the District Government of Upper Bavaria (Regierung von Oberbayern) for Animal Care (Permit Number: ROB-55.2-2532.Vet 02-17-215)

## 2. Acknowledgments

**This is a pre-print of a paper for which the official version is published in Behavioral Ecology and Sociobiology and can be accessed under doi:10.1007/s00265-021-02972-y**. We would like to thank all past members of the “Evolutionary Ecology of Variation” group of the Max Plank Institute for Ornithology, the “Behavioural Ecology” and “Evolutionary Biology” groups at the LMU, Alexander Hutfluss, Sabrina Hepner, Janna Wuelbern, all lab technicians and colleagues at the “National Reference Centre for *Borrelia*,” field assistants, and students for help in data collection.

