## Supplemental Text 1 for "Repeatable differences in exploratory behaviour predict tick infestation probability in wild great tits"

**Tick infestation probability increases with human recreational pressure and exploratory behaviour in wild great tits (*Parus major*)**

Robert E. Rollins, Alexia Mouchet, Gabriele Margos,

Volker Fingerle, Noémie S. Becker, Niels J. Dingemanse

**--Online Supplementary Material--**

**Supplemental Text 1**

We investigated whether the null effects of age and breeding density could be due to confounding effects at different levels of biological variation. For this we reran our main analysis (Table 1) with partitioning age and breeding density into two variables each. Age was split into two variables using within-subject centring (van de Pol & Wright, 2009). First, mean values for age were calculated over all observations of an individual (among-individual age as a measure of longevity), and then the deviation of a single observation from the individual’s mean was calculated (within-individual age). Density was calculated as the number of breeding pairs producing first clutches per hectare. To investigate whether temporal or spatial variation in density affected tick infestation, density was partitioned into among-plot (individual plot-mean over all the years) and among-year-within-plot variation in density (deviation within a given year from the individual plot-mean) (Nicolaus, Tinbergen, Ubels, Both, & Dingemanse, 2016). Estimates still showed no effect on infestation probability and tick burden (Supplemental Table 1) so we concluded that these effects are simply absent in our analysis.

**Supplemental Table 1.** Effect size estimates (β) and 95% credible intervals (CI) for great tit tick infestation as the probability of infestation (n = 784 records) and tick burden of infested individuals (n = 520 records).

|  | **Infestation Probability** | **Tick Burden** |
| --- | --- | --- |
| **Fixed Effects** | **β (95% CI)** | **β (95% CI)** |
| Intercept | 0.84 (0.34, 1.33) | 1.16 (0.67, 1.64) |
| Recreational pressure | 0.24 (0.02, 0.47) | -0.06 (-0.20, 0.09) |
| Among-plot variation in density | -0.17 (-0.62, 0.27) | -0.06 (-0.30, 0.19) |
| Within-plot variation in density | 0.14 (-0.11, 0.41) | 0.00 (-0.19, 0.20) |
| Exploratory score | 0.18 (0.01, 0.35) | -0.01 (-0.11, 0.10) |
| Body condition | 0.06 (-0.11, 0.23) | -0.10 (-0.20, -0.01) |
| Sex^a^ | -0.04 (-0.39, 0.30) | -0.01 (-0.20, 0.19) |
| Among-individual variation in age | -0.11 (-0.29, 0.07) | -0.05 (-0.16, 0.05) |
| Within-individual variation in age | 0.02 (-0.14, 0.19) | 0.06 (-0.01, 0.14) |
| **Random Effects** | **σ^2^ (95% CI)** | **σ^2^ (95% CI)** |
| Year | 0.05 (0.00, 0.21) | 0.16 (0.04, 0.44) |
| Plot | 0.52 (0.22, 1.00) | 0.05 (0.02, 0.10) |
| Plot-Year | 0.03 (0.03, 0.05) | 0.07 (0.04, 0.11) |
| Observer | 0.00 (0.00, 0.01) | 0.13 (0.06, 0.23) |
| Nest-box | 0.66 (0.56, 0.76) | 0.27 (0.23, 0.32) |
| Individual | 0.00 (0.00, 0.00) | 0.45 (0.40, 0.51) |
| Residual | 0.35π^2^ (0.35π^2^, 0.34π^2^) | 1.28 (1.42, 1.18) |
| ^a^Differences between sexes (reference category = female) | | |
